# Identifying Brain Region Connectivity using Steiner Minimal Tree Approximation and a Genetic Algorithm

**DOI:** 10.1101/626598

**Authors:** Syed Islam, Dewan M. Sarwar

**Affiliations:** Department of Computer Science, University College of London, London, UK; Auckland Bioengineering Institute, University of Auckland, Auckland, New Zealand

**Keywords:** Brain regions connectivity, Steiner Minimal Tree, Genetic Algorithm, SMT-Neurophysiology, SMT-Genetic

## Abstract

**Background:** Computation and visualization of connectivity among the brain regions is vital for tasks such as disease identification and drug discovery. An effective visualization can aid clinicians and biologists to perform these tasks addressing a genuine research and industrial need. In this paper, we present SMT-Neurophysiology, a web-based tool in a form of an approximation to the Steiner Minimal Tree (SMT) algorithm to search neurophysiology partonomy and connectivity graph in order to find biomedically-meaningful paths that could explain, to neurologists and neuroscientists, the mechanistic relationship, for example, among specific neurophysiological examinations. We also present SMT-Genetic, a web-based tool in a form of a Genetic Algorithm (GA) to find better paths than SMT-Neurophysiology.

**Results:** We introduce an approximation to the SMT algorithm to identify the most parsimonious connectivity among the brain regions of interest. We have implemented our algorithm as a highly interactive web application called SMT-Neurophysiology that enables such computation and visualization. It operates on brain region connectivity dataset curated from the Neuroscience Information Framework (NIF) for four species – human, monkey, rat and bird. We present two case studies on finding the most biomedically-meaningful solutions that identifies connections among a set of brain regions over a specific route. The case studies demonstrate that SMT-Neurophysiology is able to find connection among brain regions of interest. Furthermore, SMT-Neurophysiology is modular and generic in nature allowing the underlying connectivity graph to model any data on which the tool can operate. In order to find better connections among a set of brain regions than SMT-Neurophysiology, we have implemented a web-based tool in a form of a GA called SMT-Genetic. We present further three case studies where SMT-Genetic finds better connections among a set of brain regions than SMT-Neurophysiology. SMT-Genetic gives better connections because SMT-Genetic finds global optimum whereas SMT-Neurophysiology finds local optimum although execution time of SMT-Genetic is higher than SMT-Neurophysiology.

**Conclusion:** Our analysis would provide key insights to clinical investigators about potential mechanisms underlying a particular neurological disease. The web-based tools and the underlying data are useful to clinicians and scientists to understand neurological disease mechanisms; discover pharmacological or surgical targets; and design diagnostic or therapeutic clinical trials. The source codes and links to the live tools are available at https://github.com/dewancse/connected-brain-regions and https://github.com/dewancse/SMT-Genetic.

## Background

Brain systems are inherently complex. Such systems are comprised of various functional networks that enable connections among different regions of the brain. Computational models are essential to be able to discover the brain regions relationships that define the points of integration among the regions. Such relationships could be anatomical and functional [1], and can be formed into graphs [2]. Computational graph theory can be used to identify connectivity paths in such graphs, where the biological entities are represented as nodes and the relationships are represented as the edges of the graph.

Developing computational models about neurophysiology (i.e. about brain structure and function) is one of the major challenges in biomedicine. As a result of this challenge, neurologists and neuroscientists still have a number of unmet requirements in neurophysiology knowledge management. For instance, one of the key requirements is to consistently represent the relationship between (i) the results of neurophysiological examination carried out in the clinic and (ii) brain structures that are functionally responsible for the behaviour elicited by this examination. The ability to curate such information in a form of a graph where parsimonious routes can be inferred are useful to clinicians and basic scientists who want to understand neurological disease mechanisms to: a) further investigate cause and complications of a pathology, b) discover pharmacological or surgical targets to interfere with the mechanism of the disease, and c) design diagnostic or therapeutic clinical trials for a particular condition. For example, understanding brain region connectivity is vital to progress disease identification and prevention: alzheimer’s disease [3, 4], parkinson’s disease [5]; or drug discovery [6, 7].

An effective connectivity identification and visualization tool can aid clinicians and biologists to address a genuine research and industrial need. We have developed a web-based tool, SMT-Neurophysiology, that carries out calculations over a neurophysiology graph to find biomedically-meaningful paths, that could explain, to neurologists and neuroscientists, the mechanistic relationship, for example, among specific neurophysiological examinations. In particular, this tool takes into account a neurophysiology graph of the multiscale structural organization of the brain in terms of regions (nodes), and parthood and connectivity relations among these regions (edges). It enables computation and visualization of connectivity among different regions of the brain. Pharmaceutical companies can use this tool for the development of disease biomarkers (e.g. radiological image markers) or drugs that target proteins expression in particular brain regions.

SMT-Neurophysiology is able to identify and visualize the parsimonious connectivity links in a form of an approximation to the SMT [8] algorithm among a set of nodes that are of interest. SMT-Neurophysiology finds connections among a given set of brain regions via some intermediate brain regions over a neurophysiology graph. The neurophysiology graph on which our instance of the tool operates was curated from the Neuroscience Information Framework (NIF)^[1]^ [9].

However, in order to find better connections among a given set of brain regions than SMT-Neurophysiology, we have implemented a web-based tool in a form of a GA called SMT-Genetic. Here we have given priority quality of solution over speed. SMT-Genetic wins in this case because SMT-Genetic finds global optimum whereas SMT-Neurophysiology finds local optimum, although execution time of SMT-Genetic is higher than SMT-Neurophysiology. As a proof of concept, we have presented three case studies in this manuscript where SMT-Genetic finds better connections among a set of brain regions than SMT-Neurophysiology. The SMT-Neurophysiology and SMT-Genetic have been developed with highly interactive web applications that visualize brain connectivity data in a form of a graph and search for connectivity regions in response to user requests.

Prior research [10, 11] has shown that it is possible to identify the most densely area of connected brain regions. These regions are known as hubs that coordinate centrally with other regions with a view to stimulating overall brains functionality. Hubs play a vital role in making overall brain pathways and thus play high cost global network [12]. Because of such complex connectivity, brain regions form correlation by which most biomedically-meaningful relationships can be inferred from different test cases [13, 4]. A biology inspired algorithm for Steiner tree problem in networks have been proposed in [14]. By using the Steiner tree, authors optimized a physarum network. Physarum is a amoeba like tubular structure organism.

A computer simulation tool for folding RNA pathways using a GA have been proposed in [15]. The pathways are grown up based on the mutation, crossover and fitness function. This tool is a useful resource for predicting the RNA folding structure, as well as for knowing the RNA features. In [16], protein folding has been illustrated. Initially, GA starts with N number of structures and makes more organized and compact in each generation. Ideally, local regions are folded first and then moved on to the neighbours for folding. [17] presented a prediction technique of small proteins based on sequence and secondary structure using GAs. Another study in [18] investigated identification of various conformation regions in protein molecules using GA. [19] presented a GA for docking molecules of low energy conformation while binding proteins. In this case, GA could find an optimal connection among protein molecules in a large search environment.

The remainder of this paper is organized as follows: Implementation section discusses the motivation of identifying brain regions connectivity and provides a brief background of the NIF curated dataset, the Steiner tree problem and the genetic algorithm. Results section introduces SMT-Neurophysiology, SMT-Genetic, as well as presents some research questions and case studies. Discussion and Conclusions section summarize the work presented in this manuscript, as well as make concluding remarks.

## Implementation

In order to support the management, querying and articulation of data about the connectivity relationships, we have developed web-based tools that take into account: a “neurophysiology” graph of the multiscale structural organization of the brain in terms of regions (nodes), and its parthood and connectivity relations among these regions (edges). The creation of a neurophysiology graph that covers the majority of brain regions’ parthood and connectivity knowledge required the sourcing of neuroanatomical datasets from different organisms, including human, monkey, rat and bird.

The analysis of neurophysiology datasets (e.g. from clinical trials for specific disease patient populations) can elicit statistical inter-dependencies between the scores of two or more neurophysiological tests (e.g. when the score of test A increases, the scores of tests B and C decrease). Inferring connectivity links from such statistical associations provide key insights to clinical investigators about potential mechanisms underlying a particular neurological disease and which mechanisms can be targeted by specific therapies to slow or reverse the progress of brain pathology.

Our first goal was to develop a knowledge representation scheme from the NIF dataset in a form of a graph. This would enable the application of graph theory to infer hidden structures, links and dependencies. Our goal was then to allow visualization of connectivity networks using a highly interactive web application enabling easier understanding and interpretation of the underlying dataset. Functional properties of SMT-Neurophysiology is to meet these goals as follows:

1. **Data Model** – SMT-Neurophysiology should be able to operate on any undirected graph. Our instance uses a graph representing the brain regions as nodes and inter-regional relationships as edges, where edge weights are assigned based on the represented species. The graph model has the following properties:
  a. a neurophysiological test may be linked to a number of brain regions;
  b. a brain region may be bridged to another brain region via multiple routes;
  c. the size of the dataset is such that applying the multiple constraints to ensure the resulting path is biomedically-meaningful is difficult to achieve manually and reproducibly.
2. **Operations** – SMT-Neurophysiology operates on an *Input Set* (*IS*) of neurophysiological tests that correlate (e.g. tests A, B and C) in order to generate a mechanistic hypothesis of the most biologically-meaningful explanation of why the tests in the IS correlate with one another. These mechanistic hypotheses output from SMT-Neurophysiology take the form of a path over the neurophysiology graph, such that this path bridges neurophysiological test-linked brain regions over their parthood and connectivity relations. SMT-Neurophysiology takes into account explicit criteria that ensure that the resulting path offers a biomedically-meaningful hypothesis by selecting sets of edges that favour:
  a. **parsimony**: the smallest number of steps in a path,
  b. **relevance**: brain connectivity that relies predominantly on human rather than data from other organisms, and
  c. **specificity**: brain regions that are specifically associated with the clinical indices in the IS, as compared to the rest of the neurophysiological tests in the clinical mapping.

### Knowledge Representation

The brain regions dataset are curated from the NIF dataset to identify brain regions routing. Table 1 presents a sub-section of the NIF dataset and Figure 1 visualizes a sub-section of brain regions connectivity. From the dataset, we extracted brain regions and species (column B, C and D in Table 1), and then curated them into a JSON format ^[2]^ in order to construct a graph. In this graph, brain regions and the relationships are represented as nodes and edges, respectively. Specifically, edges have four attributes: first brain region, second brain region, weight and species. Weights of edges have been assigned arbitrarily as follows: 1 (homo sapiens), 2 (macaque), 5 (rat) and 7 (bird) where lower weight means higher priority, as shown in Table 2.

**Table 1.** A sub-section of brain regions Neuroscience Information Framework (NIF) dataset: column A (project/organization), B (first brain region), C (second brain region; brain regions B and C are connected), D (species), E (NIF Id for each record or database), F (Pubmed reference). We have curated column B, C and D from the NIF dataset.

| Project/Organization (A) | Brain region (B) | Brain region (C) | Species (D) | NIF ID (E) | PubMed ID (F) |
| --- | --- | --- | --- | --- | --- |
| ConnectomeWIKI | entorhinal cortex | subiculum | homo sapiens | nif-0000-24441 | - |
| ABCD | tegmentum | nucleus papillioformis | bird | nif-0000-00386 | PMID:12655512 |
| BAMS | precommissural nucleus | medullary reticular nucleus dorsal part | rat | nif-0000-00018 | PMID:10331578 |
| ConnectomeWIKI | medial entorhinal area | lateral entorhinal area | rattus norvegicus | nif-0000-24441 | - |
| BAMS | ventral posterior | visual area v4 | macaque | nif-0000-00018 | PMID:1822724 |
| BrainMaps | reticular nucleus of thalamus | ventral anterior nucleus of thalamus | macaca mulatta | nif-0000-00093 | - |
| ConnectomeWIKI | visual area 1 | visual area 2 | macaca fuscata | nif-0000-24441 | - |

**Table 2.** Subgraph statistics of brain regions dataset curated from the Neuroscience Information Framework (NIF)

| Nodes | Edges | Edge count | Out-/in-degree | Species | Edge weight |
| --- | --- | --- | --- | --- | --- |
| 1449 | 16243 | 19772 | min: 1<br>avg: 14<br>max: 196 | homo sapiens<br>macaque<br>rat<br>bird | 1<br>2<br>5<br>7 |

**Figure 1.**
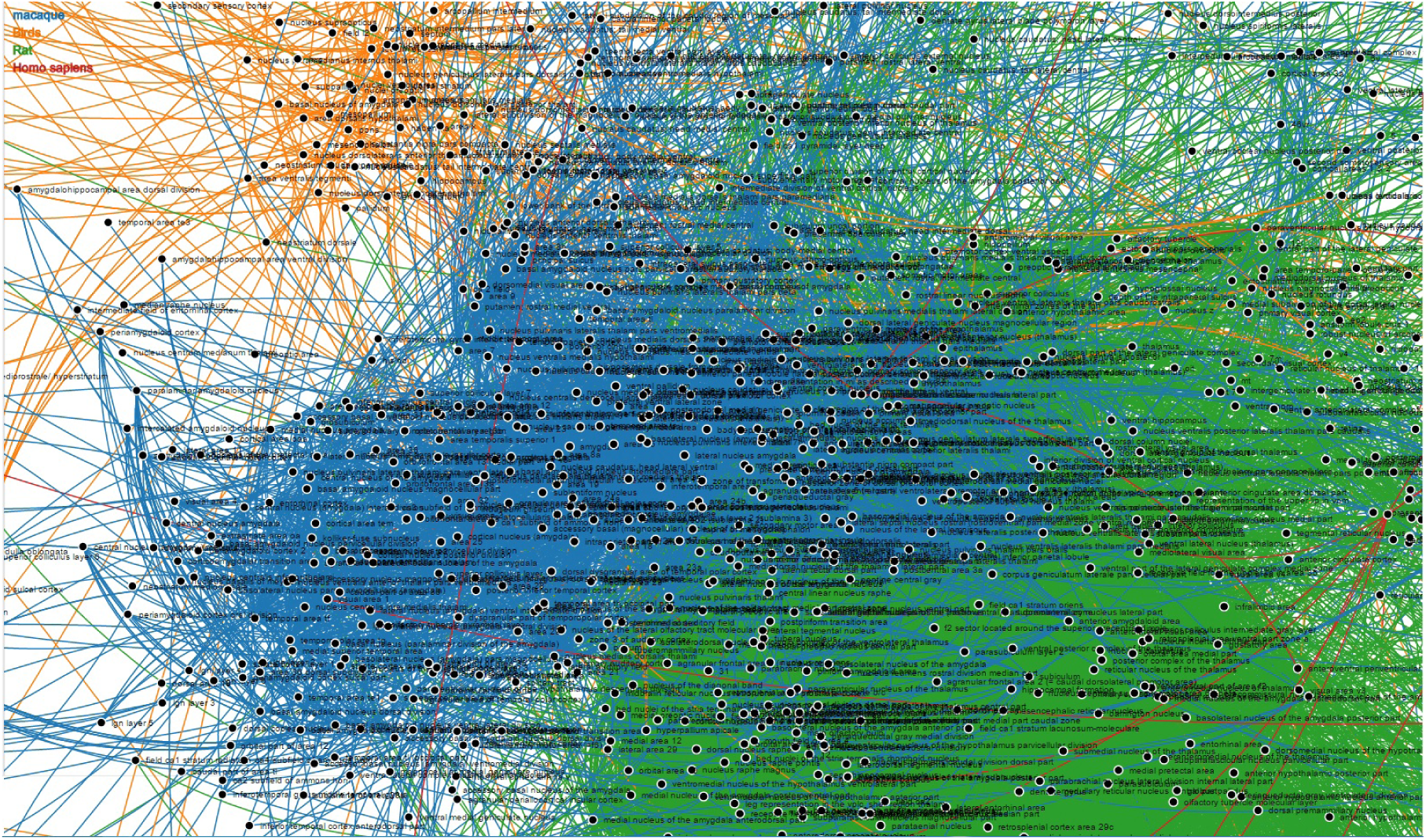
Visualization of a sub-section of brain regions connectivity curated from the Neuroscience Information Framework (NIF) dataset. On the top left corner, different colors with name of species have been used to identify biological relationships among the brain regions represented as nodes with respect to species as edges in the graph.

We have found seven species in the NIF dataset: human (homo sapiens), monkey (macaque, macaca mulatta, macaca fuscata), rat (rat, rattus norvegicus) and bird. For simplicity, we have curated four species in our JSON format by considering monkey species as macaque and rat species as rat: homo sapiens, macaque, rat and bird. The brain regions dataset consists of 1449 nodes and 16243 edges. The minimum, average and maximum outgoing edges are 1, 14 and 196, respectively presented in Table 2.

### Steiner Tree Problem

The Steiner tree [8] problem is a classical combinatorial optimization problem in graph theory. This problem is illustrated as follows: given a weighted graph with some specified vertices (i.e. terminals), idea is to find a subgraph spanning all the terminals via some intermediate vertices (i.e. Steiner points). By doing so, it tries to achieve two properties: connectivity – terminals must be connected; and optimization – find a minimum cost subgraph called minimum Steiner tree or Steiner Minimal Tree (SMT) where cost is the sum of edge weights.

### Steiner Minimal Tree

As discussed above, SMT finds connections among a set of terminals in order to achieve connectivity and optimization properties. SMT was first introduced in [20], and later SMT was discussed in a book [21] which brought attention to a large community. SMT is an NP-complete problem [22], therefore, does not guarantee an optimal solution. There are two special cases in SMT: if the terminals consist of two vertices, then SMT can be solved using Dijkstra’s shortest path algorithm [23]. Secondly, if the terminals consist of all the vertices of a graph, then SMT converges to a Minimum Spanning Tree (MST) problem. We have introduced an approximation to the SMT in order to identify the parsimonious connectivity among a set of brain regions of interest.

### Genetic Algorithm

Genetic algorithm (GA) is an evolutionary algorithm which solves search and optimization problems [24]. It is a process of iteratively recombining and reusing current instances of a problem. GA solves a problem by using some genetic concepts and operators: chromosome, population, seed, mutation, crossover, generation, and fitness function. A chromosome is a set of parameters in a form of binary or string that proposes a solution to the GA. Population is a set of chromosomes. Initial population is made from a set of chromosomes which is a seed for the GA. Execution time varies depending on the size of the chromosomes. Number of iterations is called generations. On each generation, chromosomes perform mutation – swapping bit(s) in a chromosome; crossover – recombination of bits of two parents chromosomes; and fitness function which computes a score to evaluate an optimal solution. GA terminates when a number of generations is executed. In general, the skeleton of a GA is as follows:

1. Choose an initial population of chromosomes which is a seed and evaluate the fitness function of each chromosome in the seed.
2. Crossover and Mutate chromosomes in the population.
3. Evaluate the fitness function of each chromosome in the population.
4. Select chromosomes to make a new population.
5. Exit if stopping condition or goal is met, otherwise go back to step 2 and continue.

### Greedy Algorithm

Greedy algorithm uses a heuristic to find a local optimal at each stage of a problem in order to find a global optimal solution [25]. Greedy algorithm is simple because of such computation and is useful for many applications. However, it can not always give an optimal solution. We have presented an example in Figure 5 where we applied a GA, SMT-Genetic which is a global solver, instead of our greedy algorithm, the SMT-Neurophysiology.

## Results

### SMT-Neurophysiology

We have implemented an approximation to the SMT algorithm named as SMT-approx to identify the parsimonious connectivity among a set of brain regions. SMT-approx operates on a brain region dataset curated from the NIF dataset for four species – human, monkey, rat and bird, as discussed above in the Knowledge Representation section. It finds a connection among a set of nodes called required nodes or terminals, either directly or via some intermediate nodes which are known as Steiner points. SMT-approx’s edge weights are positive integers and it is an undirected graph algorithm.

SMT-approx works as follows: each required node searches for other required nodes. When this search is successful, some of the required nodes is able to find the other required nodes, either directly or via some intermediate nodes. As such, SMT-approx forms and marks the corresponding paths as visited. These paths have been recorded as part of a solution. SMT-approx is presented in Algorithm 1. It takes nodes, edges, and required-nodes as input. Nodes is an array of brain regions and Edges is an array of four attributes: first brain region, second brain region, weight, and species. Required-nodes is an array of brain regions which must be connected either directly or via some intermediate brain regions.

SMT-approx executes N simultaneous breadth-first search (BFS) [26], one for each required node. That means, each required node maintains a separate first in first out (FIFO) queue. BFS continues until something is found or until exhaustion occurs. Exhaustion will occur when all disconnected or unsolved required nodes have empty FIFOs.

Specifically, each required node carries out 1-step of the BFS in order to search for the other required nodes unless that node is already recorded which means it is already connected or solved. For example, if an unrecorded path from a required node (n) meets another required node (m), either directly or by meeting a recorded path, then it records as part of a solution. If the meeting was direct and m was not previously recorded, then the path records m too. If an unrecorded path from a required node (n) indirectly meets another required node (m) by meeting an unrecorded path, then the union of the two unrecorded paths records both n and m unless n = m in which case then the unrecorded path has collided itself, so stop growing it. If the solution graph is not connected, it is then connected by a recursive method.

SMT-approx may not always find an optimal connection because if two required nodes meet, then the union of the two paths they take to reach that meeting point gets added to the solution and those two required nodes stop searching. If a required node does not ever meet the other required nodes, it’s search gets exhausted when it has searched its entire connected component.

We have visualized the output of the SMT-approx algorithm, SMT-Neurophysiology, in JavaScript using Data Driven Documents (D3)^[3]^ library. D3.js facilitates interactive visualization of the brain regions connectivity. Together with SMT-approx algorithm and its visualization is named as SMT-Neurophysiology. An illustrative example of SMT-Neurophysiology is shown in Figure 2. On the left panel, brain regions have been presented for the user to select. On the other hand, right panel shows visualization of the connectivity path between the selected brain regions. In order to distinguish between four species, we have used different colors on the visualization panel, as shown on the top left corner of the right panel. Red nodes represent required nodes which are selected by the user from the left panel, whereas black nodes are non-required nodes which connects the required nodes, as well as reduces the overall length of the tree.

**Figure 2.**
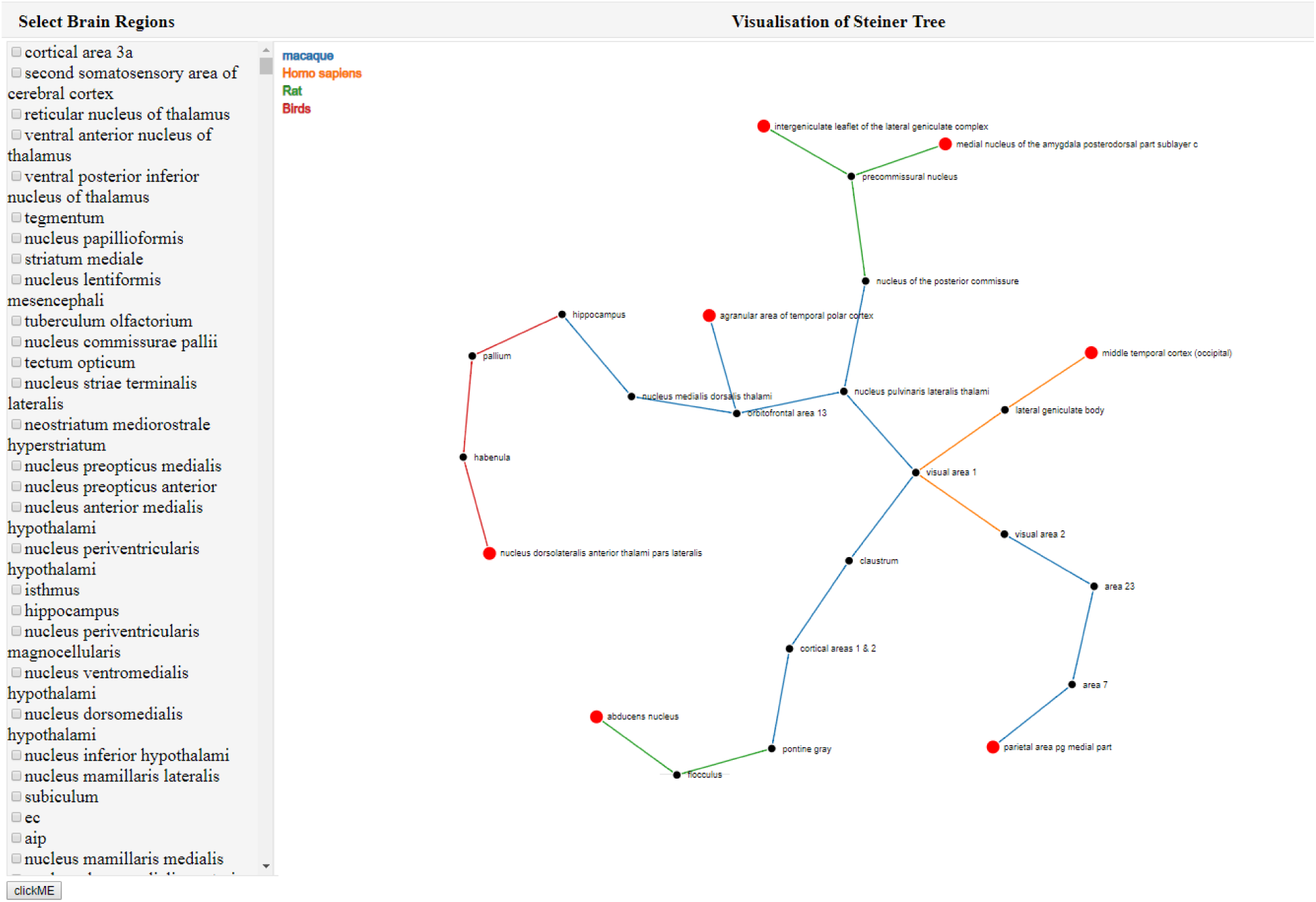
An instance of the SMT-Neurophysiology which uses the SMT approximation algorithm in the background to find connections among a set of brain regions.

Initially, SMT-Neurophysiology is rendered with left panel and an empty right panel. When the user selects some brain regions from the the left panel, then SMT-Neurophysiology executes SMT-approx algorithm in the background and finds connectivity from the curated NIF dataset.

### SMT-Neurophysiology: Case Study

The SMT-Neurophysiology finds connections among a set of brain regions, although these connections may not always be optimal because of greedy problem in nature. For this reason, SMT-Neurophysiology deviates from an optimal result. In order to address further, two research questions are presented below:

**RQ1**: *Can we identify brain regions connectivity paths using the SMT-Neurophysiology?*

**RQ2**: *What is the performance of Dijkstra*’s *shortest paths-based minimum Spanning tree to the SMT-Neurophysiology?*

RQ1: Can we identify brain regions connectivity paths using the SMT-Neurophysiology?

This research question has been investigated based on three case studies. The first two shows that SMT-Neurophysiology is able to find optimal results, whereas the third one does not find an optimal result. In order to move on to further analysis, we have assumed the following hypothesis:

#### Algorithm 1: Steiner Minimal Tree approximation algorithm

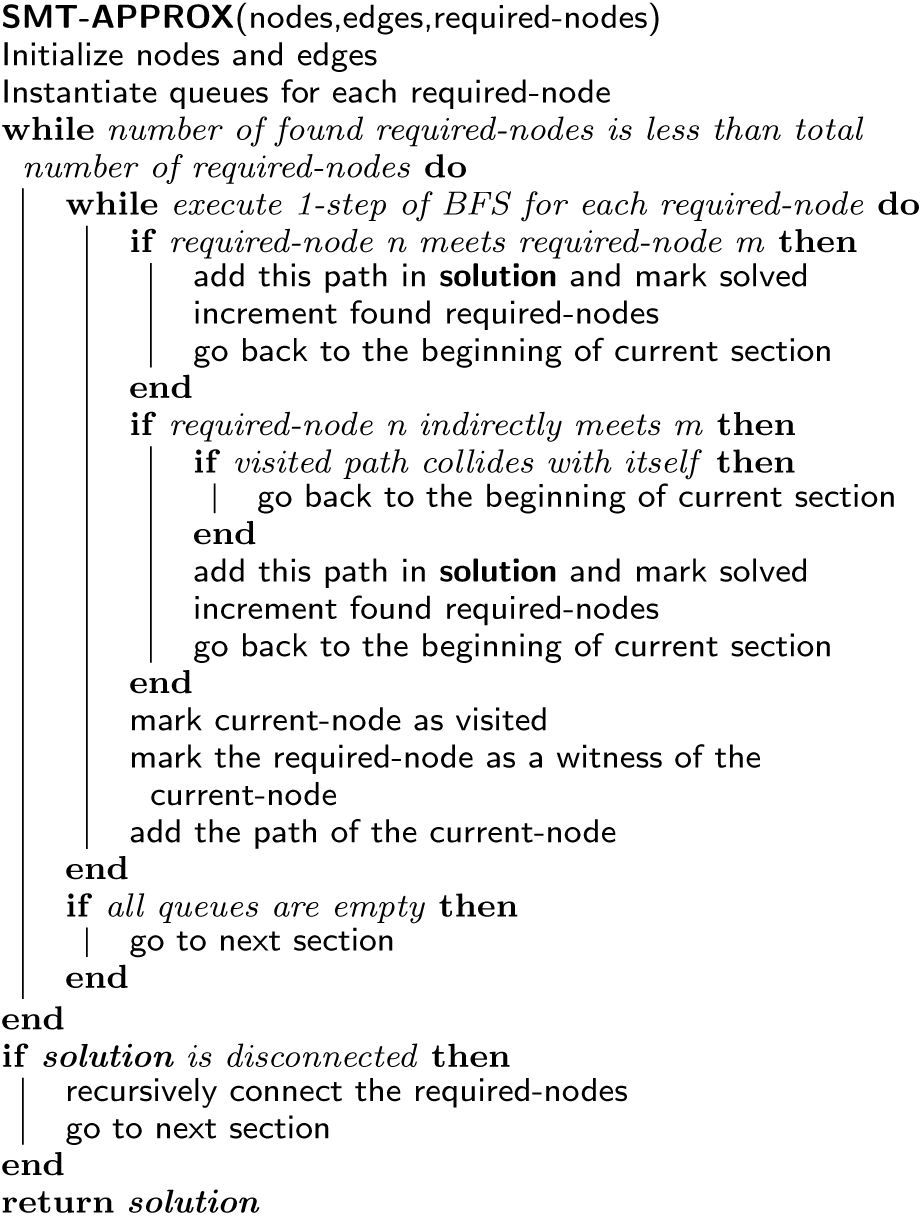

### SMT-Hypothesis

1. The Input Set (IS) consists of tests A, B and C
2. A is linked with regions x, y; B is linked with y and C is linked with w, z
3. region v is part of w
4. region v is connected to y on the basis of human data, and region z is connected to y on the basis of macaque data.

Then the most biomedically-meaningful solution that explains why tests A, B and C correlate is a mechanism over the following route: y connected to w.

### SMT-Neurophysiology: Case Study 1

Figure 3 shows a connectivity path and a correlation among a set of brain regions spanning four species by our web-based tool SMT-Neurophysiology.

**Figure 3.**
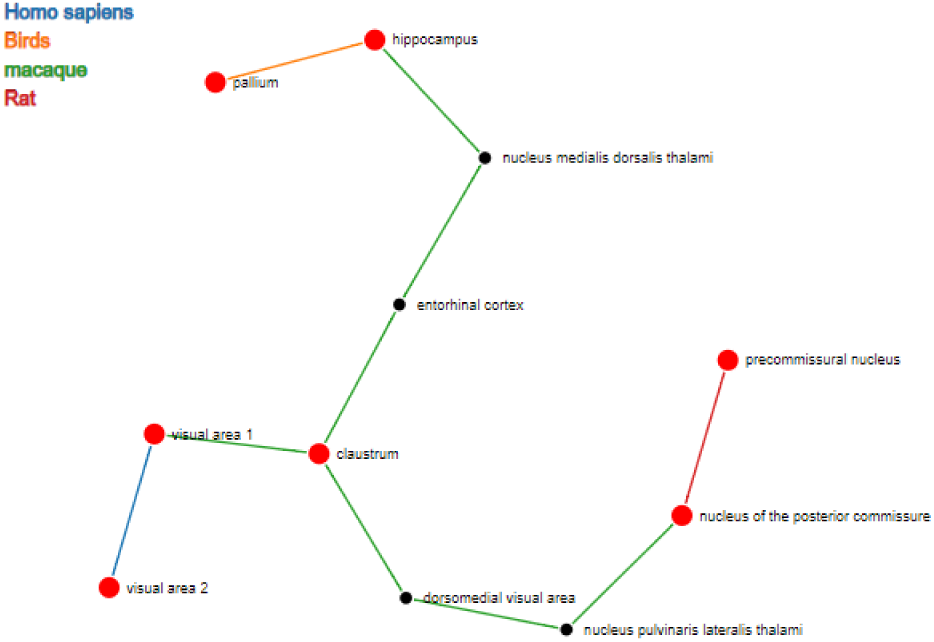
Connectivity path in SMT-Neurophysiology spanning four species – homo sapiens, macaque, rat and bird with weights 1, 2, 5 and 7 respectively.

#### Hypothesis-CS1

1. The IS consists of tests A, B, C, and D
2. A is linked with brain regions pallium, hippocampus; B is linked with hippocampus, nucleus medialis dorsalis thalami, entorhinal cortex, claustrum, visual area 1, dorsomedial visual area, nucleus pulvinaris lateralis thalami, nucleus of the posterior commissure; C is linked with nucleus of the posterior commissure, precommissural nucleus; and D is linked with visual area 1, visual area 2;
3. regions hippocampus, visual area 1, and nucleus of the posterior commissure are part of different tests;
4. region hippocampus is connected to pallium on the basis of birds data, and to nucleus medialis dorsalis thalami on the basis of macaque data. Similar explanations apply for the regions visual area 1 and nucleus of the posterior commissure.

Then the most biomedically-meaningful solution that explains why tests A, B, C, and D correlate is a mechanism over the following route: hippocampus is connected to visual area 2 and precommissural nucleus.

### SMT-Neurophysiology: Case Study 2

The second case study connects and correlates among a set of brain regions spanning three species by SMT-Neurophysiology tool, as shown in Figure 4.

**Figure 4.**
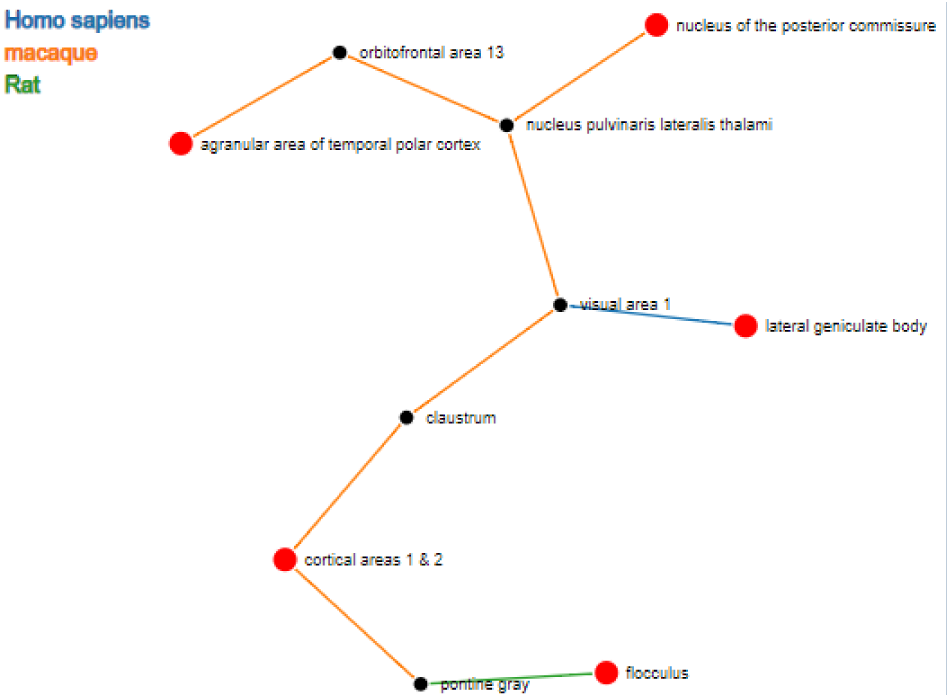
Connectivity path in SMT-Neurophysiology spanning three species – homo sapiens, macaque and rat with weights 1, 2, and 5 respectively.

#### Hypothesis-CS2

1. The IS consists of tests A, B, and C
2. A is linked with brain regions flocculus, pontine gray; B is linked with pontine gray, cortical areas 1 & 2, claustrum, visual area 1, nucleus pulvinaris lateralis thalami, nucleus of the posterior commissure, orbitofrontal area 13, agranular area of temporal polar cortex; C is linked with visual area 1, lateral geniculate body;
3. regions pontine gray, and visual area 1 are part of different tests;
4. region pontine gray is connected to flocculus on the basis of rat data, and to cortical areas 1 & 2 on the basis of macaque data. Similar explanation applies for the region visual area 1.

Then the most biomedically-meaningful solution that explains why tests A, B, and C correlate is a mechanism over the following route: pontine gray is connected to lateral geniculate body.

### SMT-Neurophysiology: Case Study 3

As discussed earlier, SMT-Neurophysiology does not always find optimal connections. Figure 5 demonstrates an example where on the left-side, a set of required nodes (red colored) make a connection via 10 macaque pathways, however, on the right-side, a better connectivity has been found via 9 pathways for the same set of required nodes. This happens because of the fact that if two required nodes meet, then the union of the two paths they take to reach that meeting point gets added to the solution and those two required nodes stop searching. If a required node does not ever meet the other required nodes, its search gets exhausted when it has searched the entire connected component.

**Figure 5.**
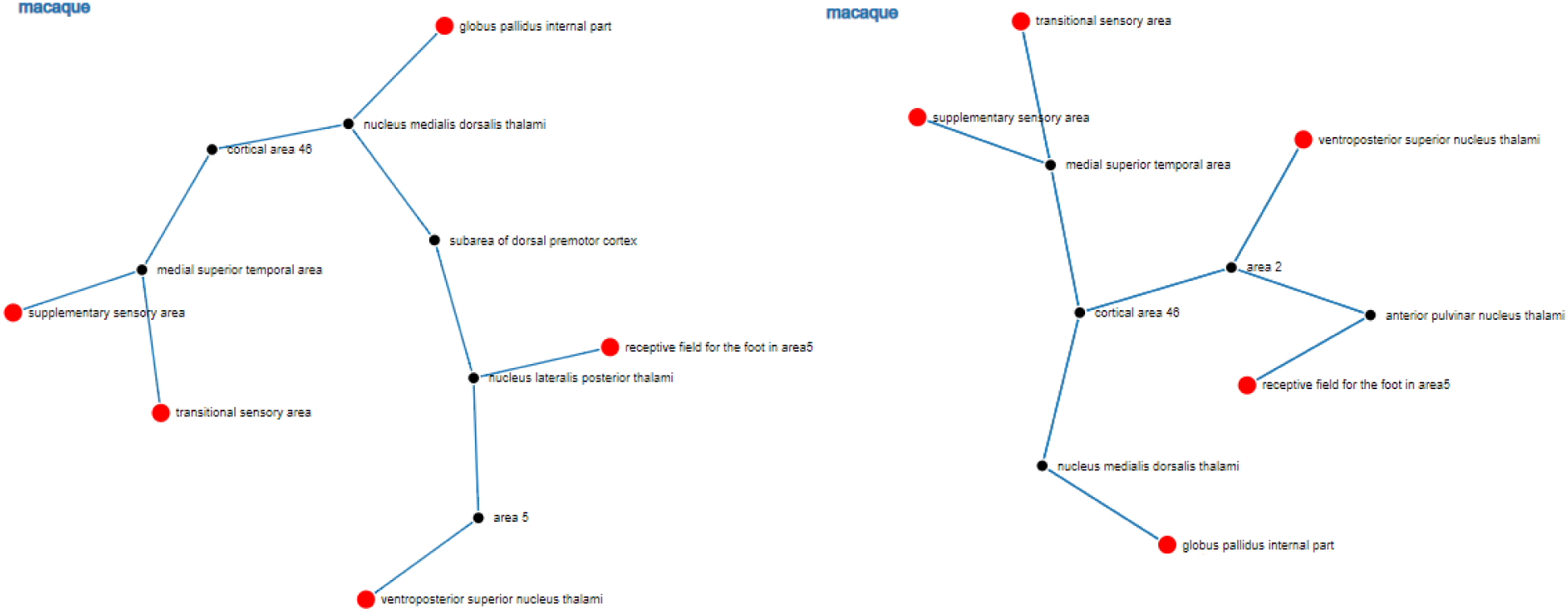
Connectivity path in SMT-Neurophysiology spanning macaque species with weight 2. On the left-side, number of edges is 10, therefore, total weight is 20, whereas on the right-side, SMT-Genetic connects 9 edges, which weight value is 18.

RQ2: What is the performance of Dijkstra’s shortest paths-based minimum Spanning tree to the SMT-Neurophysiology?

Dijkstra is a single source shortest path algorithm. In contrast, SMT-Neurophysiology finds connections among multiple sources. Therefore, shortest path strategy is not applicable to SMT-Neurophysiology. Figure 6 shows two examples. First one is the SMT of G where D is a Steiner point, and A, B, and C are required nodes. Thus, total length of the SMT of G is 12. On the other hand, second example presents G’ made of all shortest paths between the required nodes. However, SMT of G’ is not same as the SMT of G. Hence, if we consider shortest path strategy, we may not find optimal connections among a set of required nodes. For example, communication cost between B and C in the second example costs 14, whereas it only costs 8 in the first example.

**Figure 6.**
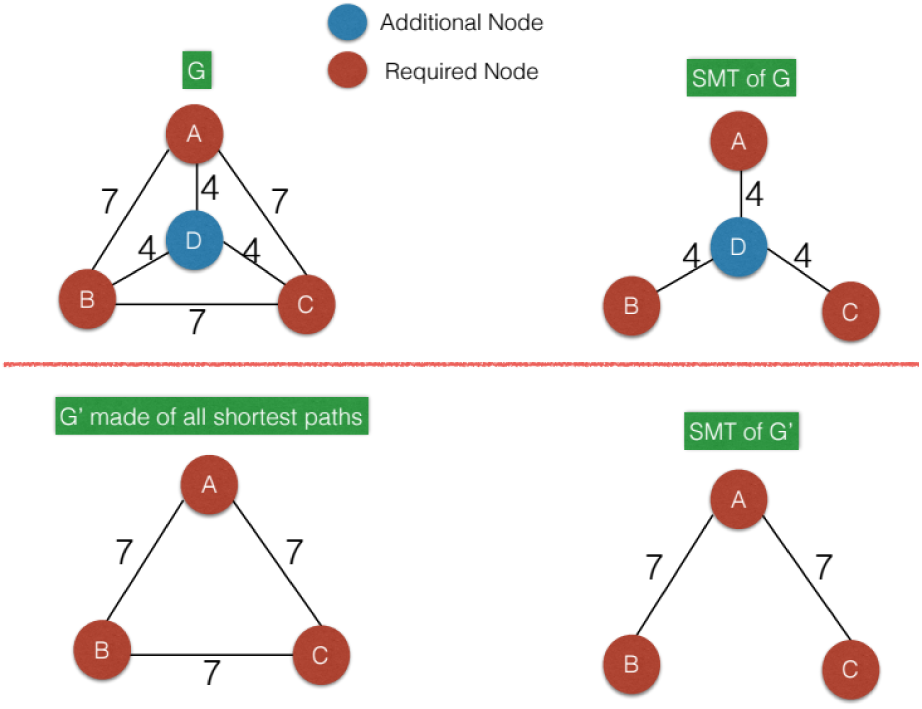
Shortest path strategy in the SMT-Neurophysiology

### SMT-Genetic

SMT-Neurophysiology uses a heuristic to find a good approximation of the SMT algorithm. Taking shortest path among all required nodes does not guarantee an optimal solution, as illustrated in Figure 6. However, GA should be able to overcome this problem by considering a longest distance between two required node of an entire SMT-Neurophysiology output graph and then expand that graph from each required node up to that longest distance. In doing so, GA will be able to find an optimal solution within the expanded graph because GA is a global solver while greedy is a local search algorithm.

In order to find better connections among a set of brain regions than SMT-Neurophysiology, we have implemented an extension in a form of a GA called SMT-Genetic. SMT-Genetic is able to find an optimal SMT-Neurophysiology for a given set of required nodes. However, SMT-Genetic deals with a large search space and thus often requires a large execution time. Various heuristics can be used to reduce such search space. To reduce the search space, we have applied the following heuristic: calculating a longest distance between two required nodes (for example, n) and then restricting the search space up to nth level by traversing through the depth-first search (DFS) [27] algorithm, which is an expansion of the SMT-Neurophysiology output. Construction of the SMT-Genetic is presented below.

### Design of Chromosomes

To design chromosomes with the NIF brain region dataset, we have set ‘1’ to represent connected edges and ‘0’ to represent disconnected edges. Figure 7 illustrates three examples of disconnected nodes’ calculation. In Figure 7 (a), node 1 is connected to node 2, and each of them has 4 disconnected nodes. Similarly, node 3, 4, 5, and 6 have 4, 4, 5, and 5 disconnected nodes, respectively. Here all nodes are required nodes and edge weight is 1. Hence, number of disconnected nodes from each required node is 26. Similar computation is done for the examples in Figures 7 (b) and 7 (c). Number of disconnected nodes is required to compute a fitness function which is discussed below. This function penalizes when the number of disconnected nodes increases; and gives preference to find a set of optimum connected nodes.

**Figure 7.**
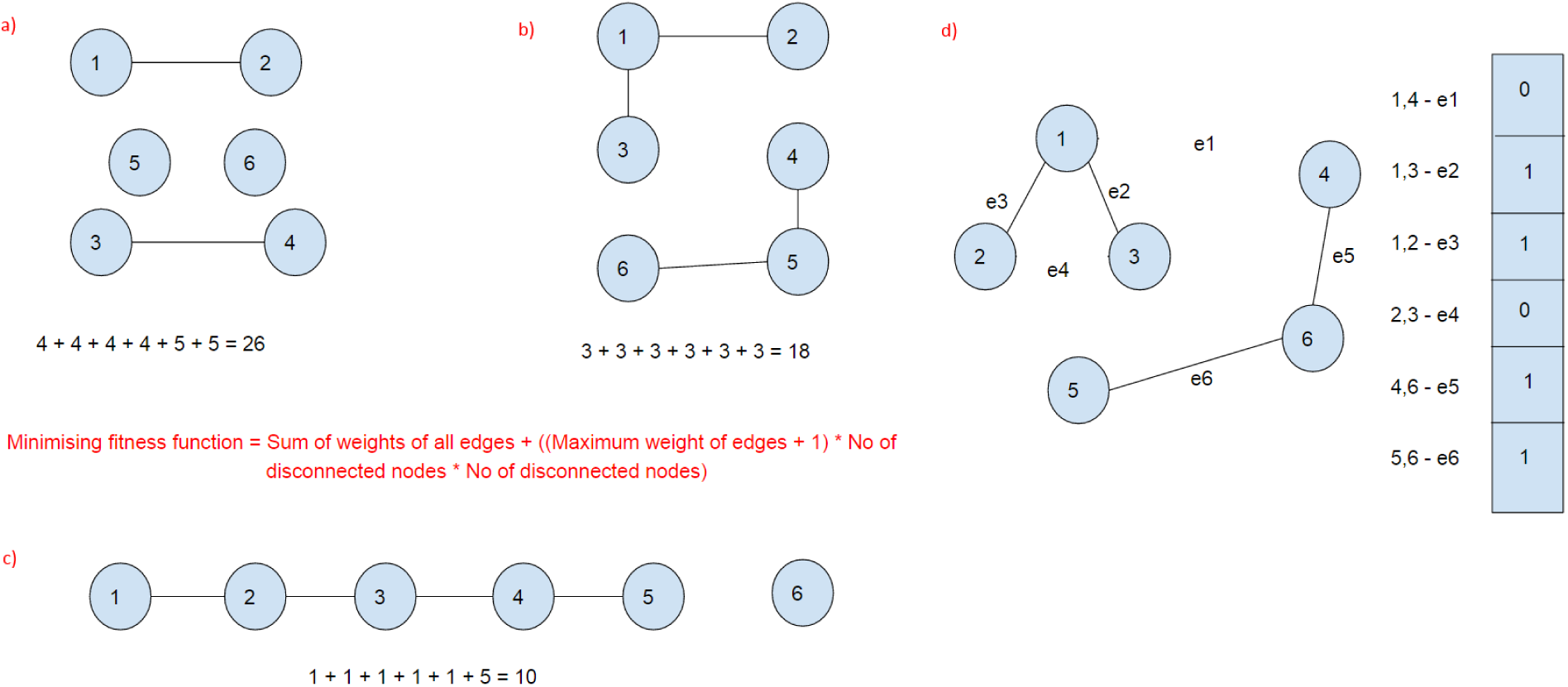
Number of disconnected nodes from each required node is in (a) 26, in (b) 18, and in (c) 10. Design of a chromosome in the SMT-Genetic is in (d) (0, 1, 1, 0, 1, 1).

A mathematical graph is created from the brain region dataset using the graph.js ^[4]^ library. The graph.js is a JavaScript library for storing arbitrary data in mathematical (di)graphs, as well as traversing and analyzing them in various ways. From each required node, we find a longest distance (weight) between the required nodes from our SMT-Neurophysiology output. Then the required nodes have been expanded from up to that longest distance using the DFS algorithm. If there is any better solution than the SMT-Neurophysiology solution within the extended graph, GA should find the better one. Our DFS algorithm is presented in Algorithm 2.

#### Algorithm 2 A DFS algorithm to extend the SMT-Neurophysiology output up to a longest weight between two required nodes

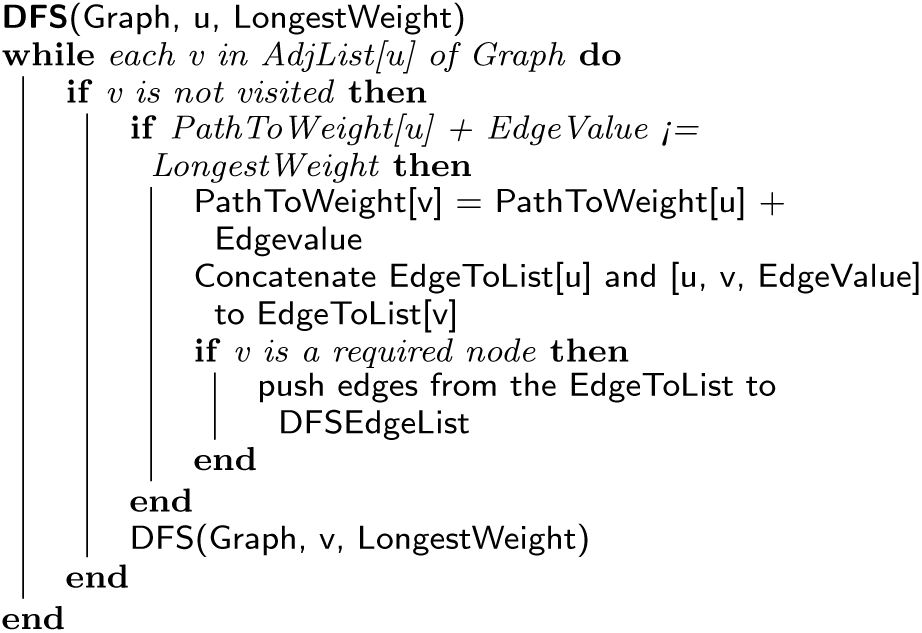

We represent edges as [node1, node2, weight] and paths as [node1, node2, …, nodeN]. Nodes, edges and required nodes, for example, in Figure 7 (c) are: [1, 2, 3, 4, 5, 6]; [[1, 2, 1], [2, 3, 1], [3, 4, 1], [4, 5, 1]]; and [1, 2, 3, 4, 5, 6]. We constructed a graph using the nodes and edges as mentioned above, and then calculated a longest distance between two required node. Here, longest distance between two required node is 4. Next, we calculated number of paths and number of edges by traversing through the entire graph in Figure 7(c) from each required node up to that longest distance.

Our DFS algorithm finds a list of paths from each required node. These paths are used to make chromosomes. In this case, each path represents a chromosome. In Figure 7 (c), for required node 1 to required nodes 2, 3, 4, 5, and 6, we get a set of paths, which is a set of chromosomes as follows: [1, 2], [1, 2, 3], [1, 2, 3, 4], [1, 2, 3, 4, 5]. From these paths, a list of edges are sorted; and the redundant edges are removed. SMT-Genetic will use these edges to find an optimum solution. Note that number of unique edges is the size of a chromosome. The above mentioned paths have the following unique edges: [1, 2, 1], [2, 3, 1], [3, 4, 1] and [4, 5, 1]. Thus the size of each chromosome is 4. Note that weight of each edge is 1. The initial chromosome is set to 0 in all indexes – [0, 0, 0, 0]. From the above paths, we split edges; and set 1 if the corresponding edges exist otherwise 0, which gives the following chromosomes: [1, 0, 0, 0], [1, 1, 0, 0], [1, 1, 1, 0] and [1, 1, 1, 1]. This calculation is shown in Table 3.

**Table 3.** Number of paths, edges, and chromosomes from each required node for the Figure in 7(c) example graph.

| Required Node | Paths | Unique edges from Paths in column 2 | Set of chromosomes |
| --- | --- | --- | --- |
| 1 | [1, 2], [1, 2, 3], [1, 2, 3, 4], [1, 2, 3, 4, 5] | [1, 2, 1], [2, 3, 1], [3, 4, 1], [4, 5, 1] | (1, 0, 0, 0), (1, 1, 0, 0), (1, 1, 1, 0), (1, 1, 1, 1) |
| 2 | [2, 3], [2, 3, 4], [2, 3, 4, 5] | [2, 3, 1], [3, 4, 1], [4, 5, 1] | (0, 1, 0, 0), (0, 1, 1, 0), (0, 1, 1, 1) |
| 3 | [3, 4], [3, 4, 5] | [3, 4, 1], [4, 5, 1] | (0, 0, 1, 0), (0, 0, 1, 1) |
| 4 | [4, 5] | [4, 5, 1] | (0, 0, 0, 1) |
| 5 | None | None | None |
| 6 | None | None | None |

If we assume that we have all the unique edges in a path, then chromosome representation of this path will be (1, 1, 1, 1). On the other hand, we have got 10 number of paths in Table 3, which is the total number of chromosomes and the size of each chromosome is 4. For example, for required node 1, paths are [1, 2], [1, 2, 3], [1, 2, 3, 4] and [1, 2, 3, 4, 5]. Path [1, 2] also represents an edge, so chromosome representation of this path is (1, 0, 0, 0). Similarly, for path [1, 2, 3], we have got two edges: [1, 2] and [2, 3]. Therefore, chromosome representation of this path is (1, 1, 0, 0).

### Seed

An initial population, seed, is required to start the SMT-Genetic. We have created 50% of the chromosomes from the combination of all 1s chromosome and the rest of them sequentially from the set of chromosomes. The population size is 250 by default, as shown in Table 4. Initial chromosome in the seed is the combination of all 1s, i.e. [1, 1, 1, 1]. A counter variable is used to keep track of the number of chromosomes. If counter is divisible by 2 then chromosomes are taken sequentially from the set of chromosomes mentioned in Table 3: [1, 0, 0, 0], [1, 1, 0, 0], [1, 1, 1, 0], [1, 1, 1, 1], [0, 1, 0, 0], [0, 1, 1, 0], [0, 1, 1, 1], [0, 0, 1, 0], [0, 0, 1, 1], [0, 0, 0, 1]. Otherwise, chromosomes are the combination of all 1s, i.e. [1, 1, 1, 1].

**Table 4.**
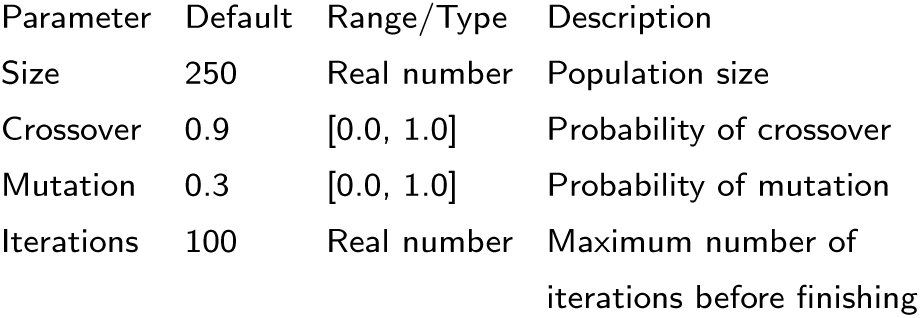
Configuration parameters in the SMT-Genetic algorithm

### Mutation

It is important to keep variation in the population to get an optimal solution. To mutate, we generate two random number and swap the corresponding bit positions of a chromosome. For example, for two random number 1 and 3, chromosome [1, 0, 0, 1] is mutated from [1, 1, 0, 0].

### Crossover

Crossover recombines bits of two parents chromosome to generate two children chromosome. We have implemented two-point crossover method where two points are randomly picked from the parents chromosome and then two children chromosome swaps the bit position of two points from their parents chromosome. We have utilized the Genetic.js ^[5]^ library to implement the two-point crossover method.

### Fitness Function

The fitness function computes a score in each iteration to evaluate an optimal solution. We have calculated minimum fitness score because our goal was to find a set of connected nodes with lower weights of edges. For this, number of disconnected nodes is more penalized than the sum of weights of all edges. However, if the graph is disconnected at a certain point of execution, maximum weight would be 0. Therefore, we have added 1 with the maximum weight of the edges. The fitness function of SMT-Genetic is presented below:

Minimizing fitness function = Sum of weights of all edges + ((Maximum weight of edges + 1) * No of disconnected nodes * No of disconnected nodes)

### Generation and Notification

Generation represents an iteration of a population, whereas notification notifies the outcome of the population. Notification is invoked after each generation. From the notification, we can find the best chromosome and can apply various statistical analysis such as summation, average, median, and standard deviation.

### Configuration parameters

The initial configuration parameters of SMT-Genetic is illustrated in Table 4. We have kept the default configuration parameters of the Genetic.js library.

Continuing with the example in Figure 7 (c), we have assigned the configuration parameters as follows: iterations: 3, size: 3, crossover: 0.9, and mutation: 0.2. Table 4 illustrates the meaning of each parameter. Number of iterations is 3, so SMT-Genetic continues up to third generation (0 to 2). For each generation, SMT-Genetic does crossover and mutation based on the assigned probability, and the fitness function checks the minimum connectivity for each chromosome among the required nodes.

### SMT-Genetic: Case Study

The research questions to be addressed in this section are:

**RQ1**: *Can SMT-Genetic find better connection when compared to SMT-Neurophysiology greedy approach?*

**RQ2**: *What is the performance trade-off in using the SMT-Genetic?*

**RQ3**: *Does it help if we seed that SMT-Genetic with the SMT-Neurophysiology greedy solution?*

RQ1: Can SMT-Genetic find better connection when compared to SMT-Neurophysiology greedy approach? SMT-Genetic can find better connection among a set of required nodes than the SMT-Neurophysiology greedy approach. Following case studies show that SMT-Genetic gives better connections among a set of brain regions than SMT-Neurophysiology.

### SMT-Genetic: Case Study 1

Figure 5 shows a connection among five required nodes where SMT-Neurophysiology connects via 10 edges (weight 20) on the left-side, and on the right-side, SMT-Genetic connects the same set of required nodes via 9 edges (weight 18).

### SMT-Genetic: Case Study 2

Figure 8 shows a connection among five required nodes where SMT-Neurophysiology connects via 10 edges (weight 20) on the left-side, and on the right-side, SMT-Genetic connects the same set of required nodes via 8 edges (weight 16).

**Figure 8.**
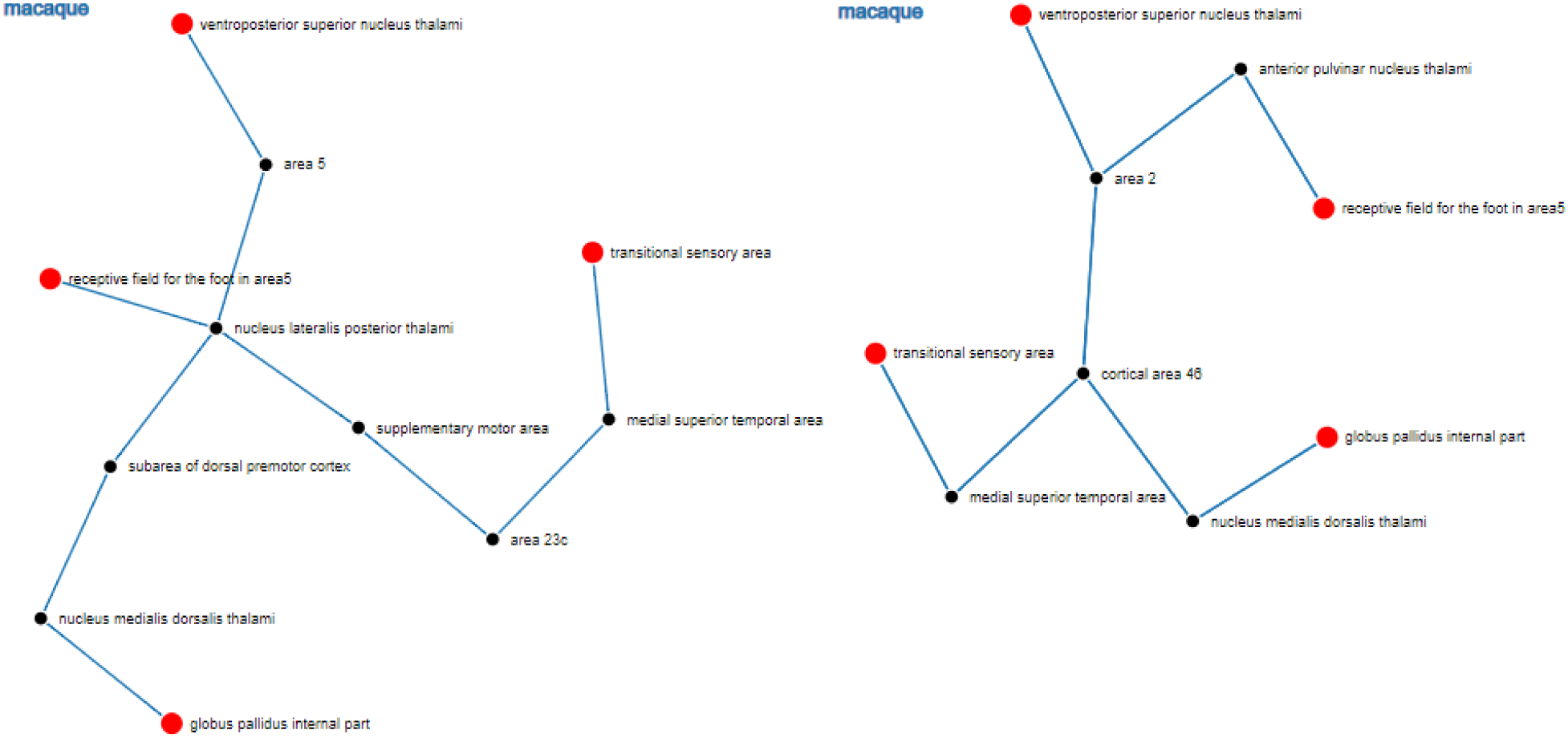
Connectivity path in SMT-Neurophysiology spanning macaque species with weight 2. On the left-side, number of edges is 10 and therefore total weight is 20, whereas on the right-side, SMT-Genetic connects with 8 edges, which weight value is 16.

### SMT-Genetic: Case Study 3

Figure 9 shows a connection among five required nodes where SMT-Neurophysiology connects via 8 edges (weight 20) on the left-side, and on the rightside, SMT-Genetic connects the same set of required nodes via 7 edges (weight 19).

**Figure 9.**
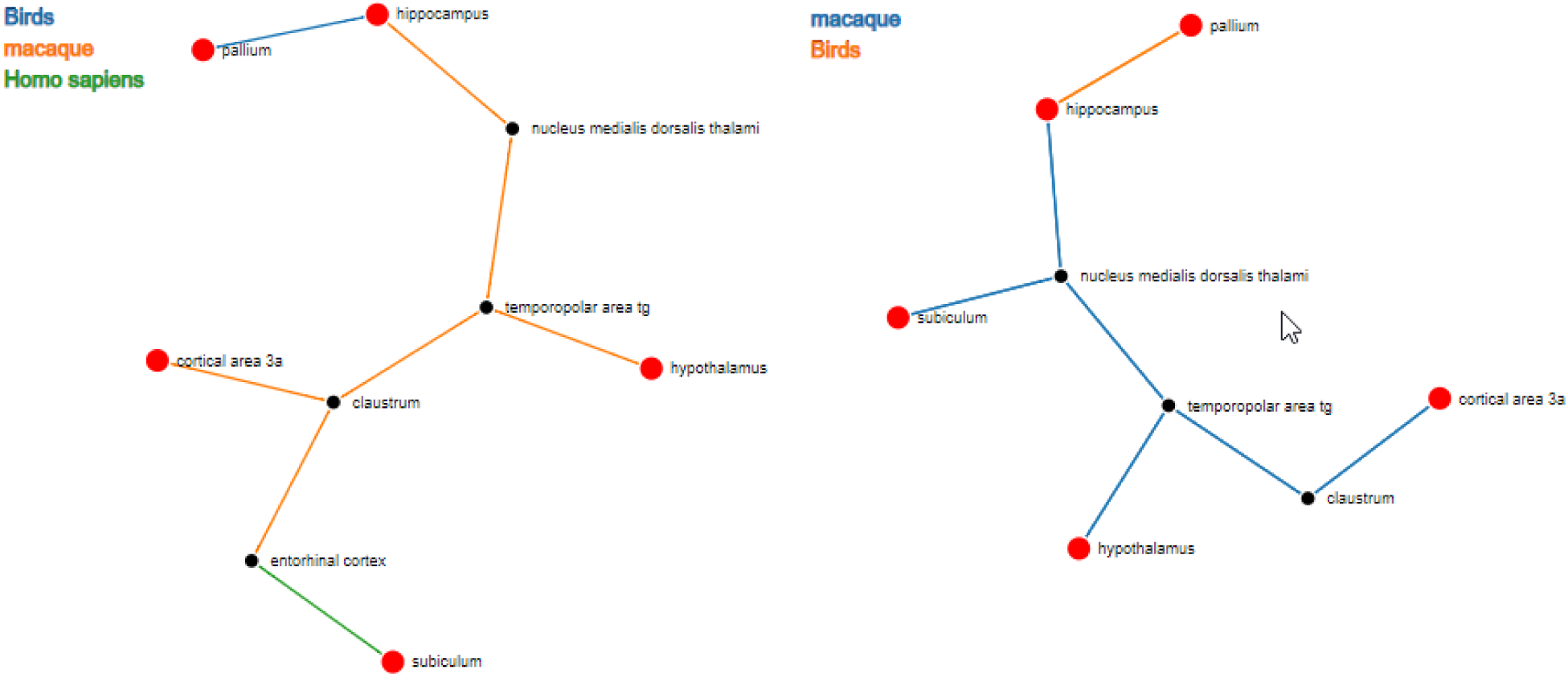
Connectivity path in SMT-Neurophysiology spanning homo sapiens, macaque, and birds species with weight 1, 2, and 7, respectively. On the left-side, weight value of 8 edges is 20, whereas on the right-side, SMT-Genetic connects with 7 edges, which weight value is 19.

RQ2: What is the performance trade-off in using the SMT-Genetic?

SMT-Genetic is able to find a better connection than SMT-Neurophysiology, as illustrated in Figures 5, 8, and 9. However, SMT-Genetic takes longer execution time. In each generation, SMT-Genetic looks for a better solution with the help of fitness function’s score. In some cases, SMT-Genetic finds an optimal solution and repeats the same solution till the end of the number of generation. In that case, we could stop the execution if SMT-Genetic repeats the same solution for a certain number of generation. For example, if a solution is repeated more than 50 times, SMT-Genetic will terminate.

RQ3: Does it help if we seed that SMT-Genetic with the SMT-Neurophysiology greedy solution?

The SMT-Genetic approach uses a DFS algorithm to traverse the brain regions dataset up to a longest distance between two required node of a SMT-Neurophysiology solution. If there exists any better solution than the SMT-Neurophysiology solution in the refined search space, SMT-Genetic should find the better one. The DFS algorithm is presented in Algorithm 2.

## Discussion and Conclusions

In this paper, we have introduced an approximation to the SMT algorithm to identify the most parsimonious connectivity among the brain regions of interest. We have implemented our algorithm as a highly interactive web application called SMT-Neurophysiology that enables such computation and visualization. It operates on brain regions connectivity dataset curated from the NIF dataset for four species – human, monkey, rat and bird.

We have presented two case studies on finding the most biomedically-meaningful solutions that identifies connections among a set of brain regions over a specific route. The case studies demonstrate that SMT-Neurophysiology is able to identify several interesting pathways among brain regions of interest. Furthermore, SMT-Neurophysiology is modular and generic in nature allowing the underlying connectivity graph to model any data on which the tool can operate. In order to find better pathways than SMT-Neurophysiology, we have implemented an extension in the form of a GA called SMT-Genetic. We have further presented three case studies where SMT-Genetic finds better connections among a set of brain regions than SMT-Neurophysiology.

Our analysis would provide key insights to clinical investigators about potential mechanisms underlying a particular neurological disease. The tools and the underlying data are useful to clinicians and scientists to understand neurological disease mechanisms; discover pharmacological or surgical targets; and design diagnostic or therapeutic clinical trials. Pharmaceutical companies can use these tools for the development of disease biomarkers (e.g. radiological image markers) or drugs that target proteins expression in particular brain regions.

In future, we want to use SMT-Genetic to achieve better results as opposed to the SMT-Neurophysiology greedy approach. Evolutionary and optimization algorithms will be used to address the greedy problem of the SMT-Neurophysiology. These search algorithms by their nature deal with a large search space and often require extremely large amount of time to run. Various heuristics could be used to reduce the search space in order to reduce the resources and run-time requirements.

To improve the performance of the SMT-Genetic by combining global and local search, we would like to propose some thoughts as follows: If we operate under the assumption that while different instances of the problem will get different required nodes, most if not all instances will run on the same graph. Under this assumption we can make run-time queries really fast if we spend significant time beforehand to optimize for a specific graph we intend to use and perform pre-processing. The pre-processing step will identify the best search strategy to retrieve SMT in (near) linear time which when employed will give the feeling of interactive search results for the end-user.

A promising technique that may be used to achieve this pre-processing is a neural network trained with a deep learning algorithm. The pre-processing phase will take a long time to run (may be days), but only has to run once, and will find a near-optimal search strategy for the run-time algorithm. The expectation is to have near linear-time speeds for run-time queries. Note that if small changes to the underlying graph are made, it may not be necessary to start the preprocessing phase all over. Training the neural network may be done incrementally.

## Declarations

### Acknowledgements

The authors would like to thank the Open Physiology research group for valuable discussions and feedback for making this work possible. Authors are grateful to the Medical Technologies Center of Research Excellence (MedTech CoRE), the Aotearoa Foundation, and the Auckland Bioengineering Institute for supporting this project.

## Availability of data and materials

The source codes and links to the live tools are available at https://github.com/dewancse/connected-brain-regions and https://github.com/dewancse/SMT-Genetic.

## Authors’ contributions

Dewan developed SMT-Neurophysiology visulization application. Syed and Dewan designed and developed SMT-Genetic application. Bernard analyzed the case studies. Dewan and Syed wrote and revised the paper. All authors read and approved the final manuscript.

## Footnotes

[1] www.neuinfo.org

[2] https://raw.githubusercontent.com/dewancse/SMT-Genetic/master/data.json

[3] https://d3js.org/

[4] https://github.com/mhelvens/graph.js

[5] http://subprotocol.com/system/genetic-js.html

